# A structural database of chain-chain and domain-domain interfaces of proteins

**DOI:** 10.1101/2022.04.13.488238

**Authors:** Neeladri Sen, M.S. Madhusudhan

**Author notes:** To whom the correspondence should be made.

## Abstract

In this study, we have mined the PDB and created a structural library of 178,465 interfaces that mediate protein-protein or domain-domain interactions. Interfaces involving the same CATH fold(s) were clustered together. Our analysis of the entries in the library reveals the similarity between chain-chain and domain-domain interactions. The library also illustrates how a single protein fold can interact with multiple folds using similar interfaces. The library is hence a useful resource to study the types of interactions between protein folds. Analyzing the data in the library reveals various interesting aspects of protein-protein and domain-domain interactions such as how proteins belonging to folds that interact with many other folds also have high EC values. These data could be utilized to seek potential binding partners. It can also be utilized to investigate the different ways in which two or more folds interact with one another structurally. We constructed a statistical potential of pair preferences of amino acids across the interface for chain-chain and domain-domain interactions separately. They are quite similar further lending credence to the notion that domain-domain interfaces could be used to study chain-chain interactions. Lastly and importantly, the library includes predicted small molecule binding sites at the protein-protein interfaces. This has applications as interfaces containing small molecule binding sites can be easily targeted to prevent the interaction and perhaps form a part of a therapeutic strategy.

## 1. Introduction

Protein-protein interactions are vital for several biological processes [1,2]. Identifying and characterizing such interactions could help explain the functioning of proteins and the basis of various diseases [3,4]. Various databases such as Database of Interacting Proteins (DIP) [5], Biomolecular Interaction Network Database (BIND) [6,7], Molecular Interaction Database (MINT) [8], Interactome3D [9] etc. list experimentally validated interactions. Except for Interactome3D none of the other lists contains structural information.

The 3D structure of complexes of interacting proteins help explain the mechanism of interaction, which in turn shed light on the functioning of cellular pathways [10–13]. 3D structures of these complexes can be determined by X-ray crystallography, NMR spectroscopy, cryo-EM, etc. Though the number of 3D structures of proteins complexes is steadily increasing, these experiments are expensive, labor intensive and technically challenging [14,15], necessitating computational approaches [15–21]. With the advent of high accuracy deep learning driven methods such as AlphaFold [22] and RoseTTAFold [23] for predicting the 3D structures of proteins, the next frontier is to make accurate predictions of the structures of protein complexes. One of the key ingredients of the success of these prediction methods was the availability of over 100,000 structures and over 200 million sequences of proteins. There are however far fewer protein complexes in the PDB (of the order of thousands). The same deep learning techniques have been adapted for the prediction of the structures of protein complexes [24,25]. However, to improve these techniques, we would need to either wait for more data on 3D structures of complexes to accumulate or leverage information from known structures, paying particular attention to domain-domain interfaces. Our study, which primarily focuses on binary interactions, is an effort in the latter direction. An argument can be made that protein cores and protein interfaces show similar physico-chemical properties (such as amino acid composition, contact preferences etc) and so monomeric structures could help build structures of complexes. However, cores and interfaces differ in structural packing and composition [26]. Hence the vast repertoire of protein structures that are not in complexes cannot be used to model protein complexes and we need to rely on data from protein interfaces.

Large proteins contain multiple domains, which are defined as independent folding, evolving and structural units in proteins. Two proteins chains can fuse (gene fusion) leading to the formation of two protein domains in a single protein. Conversely, two (or more) domains of the same protein can split and evolve into two (or more) independent chains [27,28]. During these fusion/fission events, a chain-chain interface can convert to a domain-domain interface or vice versa. Hence interfaces on domains can be structurally similar to that between chains. The domain definitions/boundaries of individual proteins have been characterized in SCOP/SCOPe [29,30], CATH [31–33], Pfam [34] and Ecod [35] which can be utilized to identify interfaces between different protein domains.

Multiple libraries have been developed in the past to characterize protein-protein/domain-domain interfaces such as 3DID [36–38], PIBASE [39], SCOPPI [40], SNAPPI-DB [41], SCOWLP [42,43], ProtCID [44]. Many of these databases classify and cluster the interfaces to show similarities between different interfaces or study specific properties of the interfaces such as conservation, the importance of water etc. Methods such as PRISM [16] and InterComp [45] utilizes the interfaces as templates to model protein complexes.

Protein complexes can be modeled either by docking one protein onto another or by comparative modeling using a templates protein complex. Template based modeling of protein complexes has been shown to be more accurate in comparison to docking [46– 48]. In addition, recent literature indicates that the structural repertoire of protein interfaces is degenerate and close to complete [49,50] and nature reuses similar interfaces across different proteins. Hence a library of such observed protein-protein/domain-domain interfaces will be useful in understanding and modeling protein complexes. A combined domain-domain and chain-chain interface library might be useful and may account for gene fusion/fission events. Hence, a composite library might provide a better sampling of the structural space of the protein interfaces.

We have created a library of all known interfaces between different proteins (chain-chain interface) and also separately catalogued intra-chain domain-domain interfaces. Libraries as such can be useful in studying how protein folds structurally interact with one another. Thus, we structurally clustered interfaces belonging to the same fold, to identify the various modes of interactions between proteins belonging to the same fold. Using the interface library, we showed how domain-domain and chain-chain interfaces and non-homologous protein complexes (belonging to the same/different folds) could have structurally similar interfaces. In addition to structurally characterizing the interfaces, we compared the amino-acid pair preferences between domain-domain and chain-chain interfaces. We also showed that the interfaces are more conserved as compared to the whole protein. The library also contains predicted small molecule binding sites that could be targeted to prevent protein complex formation, with possible therapeutic applications.

Regardless of how proteins interact, be it homologous pairs interacting similarly [51,52] or unrelated protein pairs (or proteins and peptides) using similar interfaces [45] [49], our interface library could be used to predict the 3D structures of the complexes.

## 2. Results

### 2.1. Database of interfaces

We extracted 112,043 binary chain-chain interfaces and 66,442 binary domain-domain interfaces (domain definitions based on CATHv4.2) from 42,259 PDB structures (table 1). As mentioned in the methods section, we call these fold combinations. Note that only 89,993 out of 112,043 fold combinations have an associated CATH identifier. This has implications on how these fold combinations are clustered (see section 2.3). The 22,110-fold combinations that are not annotated by CATH ids come from 6712 PDB entries. These interfaces are still a part of the library, albeit without being clustered (see section 2.3).

**Table 1.**
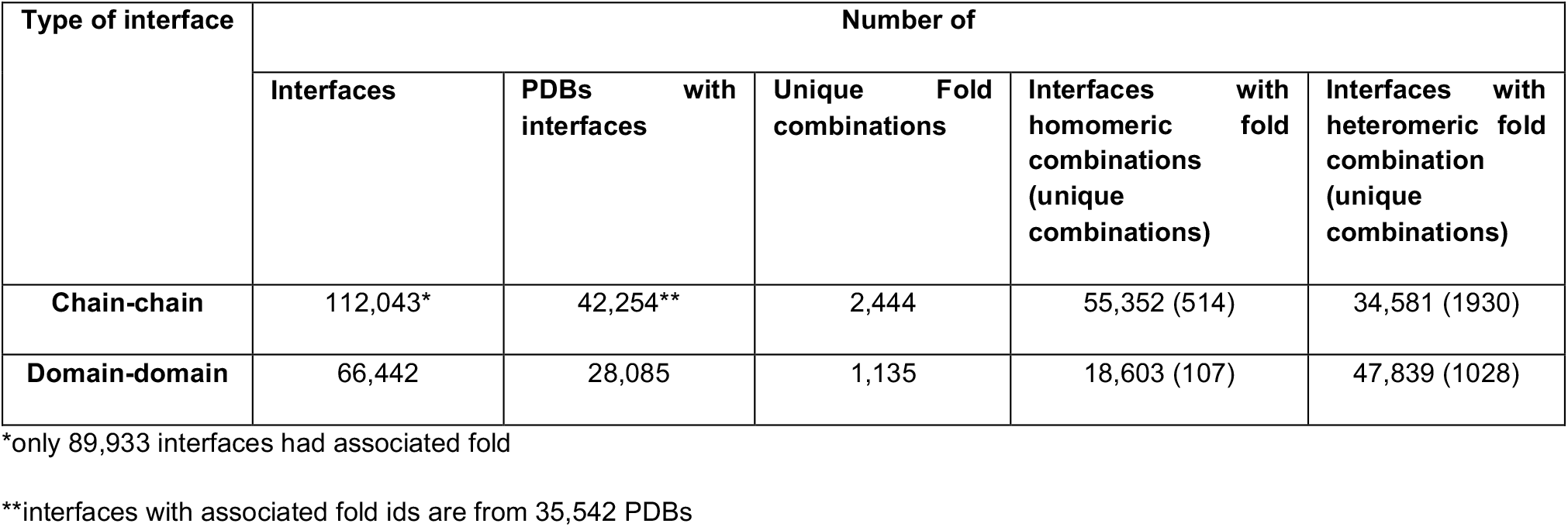
Composition of the interface library PDBs.

Binary interfaces are made of both homo- and heteromeric interactions. Among chain-chain interfaces, 62% of fold combinations were homomers, whereas this was only 28% of the domain-domain interfaces. Homomeric associations are more abundant in interactions between protein chains than interactions between protein domains.

### 2.2. Fold related properties of the interface library

#### 2.2.1. Fold Combination

The binary fold combinations in our library contain records of CATH ids interacting with one another. There are 524 CATH ids that interact with only one other fold and on the other extreme, the CATH id 3.40.50 (Rossman fold) interacts with 238 other folds. The number of fold combinations that a CATH id is a part of has a correlation coefficient of 0.82 with the number of its Enzyme Commission (EC) terms (Figure S1). Generally, the higher the number of EC terms associated with a fold, the greater the number of folds it would interact with. Of the 1391 CATH folds, only 1109 folds interact with one another. Of these, 1092 folds interact with <30 other folds. The Rossman fold (CATH ID-3.40.50) has the highest number of fold combinations, 238, to go along with over 1,000 EC terms (Table2). CATH id 1.20.5 (Single alpha-helices interacting in coiled-coils or other helix-helix interfaces) interacts with 90 different folds, perhaps because of the diversity of the sequences that can take up this particular fold [53]. The list of 15 folds that are a part of >30 fold combinations (table 2) contains some of the most abundantly populated CATH ids. The fact that some folds such as Rossmann fold, TIM barrels and Jelly rolls have more EC terms (1087, 317 and 132 respectively) than they appear in fold combinations (238, 66 and 57 respectively) could imply that several fold combinations of these folds are yet to structurally explored. The higher the number of EC terms in a fold, the more varied the functions of the proteins in the fold, hence the larger the number of folds they interact with.

**Table 2.** Folds that participate in >30 binary fold combinations.

| CATH ID | Fold Name | Number of fold combinations | Number of EC terms per fold |
| --- | --- | --- | --- |
| 3.40.50 | Rossmann fold | 238 | 1087 |
| 2.60.40 | Immunoglobulin fold | 93 | 111 |
| 1.20.5 | Single alpha helix involved in coiled-coil or other helix-helix interactions | 90 | 22 |
| 3.30.70 | Alpha-beta plaits | 81 | 124 |
| 1.10.287 | Helix hairpins | 79 | 45 |
| 1.10.10 | Arc-repressor mutants, subunit A | 78 | 61 |
| 3.20.20 | TIM barrels | 66 | 317 |
| 2.60.120 | Jelly rolls | 57 | 132 |
| 1.25.40 | Ser-Thr protein phosphatase 5, Tetratricopeptide repeat | 54 | 54 |
| 1.20.58 | Methane monooxygenase hydrolase Chain G, Domain 1 | 54 | 16 |
| 2.40.50 | OB-fold (Dihydrolipoamide acetyltransferase. E2P) | 48 | 56 |
| 3.10.20 | Ubiquitin like rolls | 38 | 49 |
| 3.30.420 | Nucleotidyl transferase domain 5 | 37 | 48 |
| 2.30.30 | SH3 type barrel | 37 | 36 |

#### 2.2.2. Number of interfaces per fold combination

The 1109 folds that form chain-chain and domain-domain interfaces feature in 3,065 unique fold combinations. 585 fold combinations have only one known interface (1 unique chain-chain or domain-domain interface in 1 fold combination). 2,502 (82%) fold combinations had <=30 interfaces (Figure S2). An interface here refers to a particular configuration of a fold combination. A single fold combination could be observed in the PDB to have multiple interfaces. A homo-oligomeric structure could have various instances of the same interface within a single PDB entry.

Homo-dimeric interactions between Rossmann folds (14,844 interfaces), Immunoglobulin-like folds (6,528 interfaces) and Glutamine Phosphoribosylpyrophosphate-subunit 1-domain 1 folds (6,318 interfaces) are the three largest populations of fold combinations. As was shown earlier (Table 2), the Rossmann fold and the Immunoglobulin like fold are associated with 1,087 and 111 EC Terms respectively. In addition, the Rossman Fold is the most diversified and prevalent fold of ancient evolutionary origin and accounts for about 15% of the human proteome [54]. A large number of instances (6,318) of Glutamine Phosphoribosylpyrophosphate-subunit-1-domain-1 fold homo-dimers are found in only 277 PDB entries, as most/all of these entries contain homo-multimers of the same protein. This fold too has 248 EC terms. This indicates the diversity in the sequence and functions of proteins adopting these folds.

### 2.3. Clustering of the interface library

Given the number of instances the same fold combinations appear in different interfaces, we clustered (within each fold combination) the 156,375 interfaces ((89,933 chain-chain and 66,442 domain-domain) according to structural similarity. To find general structural patterns in the ways folds and domains interact with one another, we grouped them into structural clusters within each fold combination. This clustering can also help identify if non-homologous protein pairs can use the same interface geometry and if chain-chain and domain-domain interfaces are structurally similar. The cluster representatives can also serve as structurally non-redundant templates for the comparative modelling of protein complexes or multidomain proteins. The interfaces were grouped into 27,885 clusters that have structure overlap >=80% and RMSD <=1.5 Å with respect to that of the representative PDB. 13,039 (∼47%) clusters only contain 1 PDB (Figure S3), which might be a result of the stringent RMSD and structure overlap criterion used during clustering. 26,401 (∼95%) clusters contain less than 20 PDB per cluster (Figure S3).

#### Superfamily based investigation of interface clusters

Each CATH fold is further subdivided into homologous superfamily. We checked if the structural clusters formed as described above were within a superfamily. Out of the 27,885 clusters, 27,741 clusters (99.5%) had all interfaces belonging to the same superfamily combination. Only 144 clusters had interfaces belonging to different superfamily combinations. Out of these 144 clusters, 130 clusters had interfaces from a combination of 2 superfamilies. Of the remaining 14 clusters having more than 2 superfamily combination in the same cluster, 9 clusters belonged to the fold 1.20.5 (Single alpha helix involved in coiled-coil or other helix-helix interactions). This is because the single alpha helix fold is a secondary structural component that could easily interact with other helices from various superfamilies, leading to the clustering together of multiple superfamilies. Even though our study was done at a fold level, 99.5% of the clusters were formed superfamily wise, indicating that interfaces are structurally conserved within superfamilies. Had we used less stringent criteria as cut-offs for clustering, we might have had more clusters across superfamilies.

#### 2.3.1. Number of interface clusters per fold combinations

The number of clusters per fold combination ranges from 1 to 8,004. 1,626 fold combinations (out of the 3,065 observed combinations) had only 1 cluster (585 fold combinations had only 1 interface and hence only had 1 cluster). 2,942 (96%) fold combinations had <20 clusters (Figure S4). 3043 (99%) fold combinations were clustered into less than 100 clusters.

The three highest numbers of self-interactions are those involving the Rossmann fold, immunoglobulin-like fold and TIM barrel, with 14,844, 6,528 and 3,598 interfaces respectively. These interfaces were grouped into 7,809, 1,675 and 984 clusters respectively. The high number of clusters in these fold combinations was because multiple non-homologous proteins take up these folds [55] (as indicated by high EC numbers for the fold - Table 2). This results in multiple modes of interactions depending on the protein type. Most of the interfaces (73%) of self-interactions between Rossman folds have high RMSD (>2 Å) and low structure overlap (<70%) when the different protein interfaces are structurally superimposed on the cluster representatives (Figure S5).

In certain cases, we have split PDB entries belonging to the same fold combination into multiple clusters because of the stringency in our clustering criterion. Certain folds such as the hemagglutinin-ectodomain chain B had 484 out of 567 interfaces clustered together with cluster representative 4gxx_BD. The other interfaces were grouped into different clusters because they had a structural overlap ranging between 70-80% and RMSD between 1.5 Å – 2.4 Å with 4gxx_BD and hence were not clustered together. It is clear that the stringency of clustering modulates the number of clusters. In addition, these stringent clusters can help in modeling protein interfaces by providing better resolved clusters (Figure S6).

#### 2.3.2. Similarity between domain-domain and chain-chain interfaces

Over the course of evolution, protein domains can split into chains or protein chains can come together to form domains. The interface library could be used to find evidence of structurally similar chain-chain and domain-domain interfaces. Hence, a composite library such as the one described here can serve as a source of templates to model both protein complexes and multidomain proteins.

514 fold combinations contain both chain-chain and domain-domain interfaces. From these fold combinations, 102 clusters contain both chain-chain and domain-domain interfaces clustered together (Supplementary Text 1). These clusters had a median of 57% of domain-domain interface and 43% chain-chain interface. Clusters, as such, highlight the fact that nature reuses the same geometry across different types of interfaces.

Here are a few interesting examples that illustrate the usage of the same interface in domain-domain and chain-chain interactions, even when the proteins with those folds/domains are unrelated to one another. The domain-domain interface of Giardia dicer is superimposed onto a chain-chain interface of the Nuclease domain of ribonuclease 3; with a structure overlap of 86% and an RMSD of 1.33 Å (Figure S7A). The two proteins share a sequence identity of 23% and belong to the Ribonuclease iii N terminal endonuclease domain, Chain A fold. In the other example, the chain-chain interface of AVA_4353 protein superimposes onto the domain-domain interface of PhuS protein with a structure overlap of 91% and RMSD of 1.28 Å (Figure S7B). The two proteins belong to the heme utilizing iron like fold and share no significant sequence similarity. Curiously, within a complex of a protein carboxysome shell protein CcmP (with two domains of the same fold Alpha Beta Plaits), we see structurally similar chain-chain and domain-domain interface (Figure 1). The two domains within a monomer however share no significant sequence similarity.

**Figure 1.**
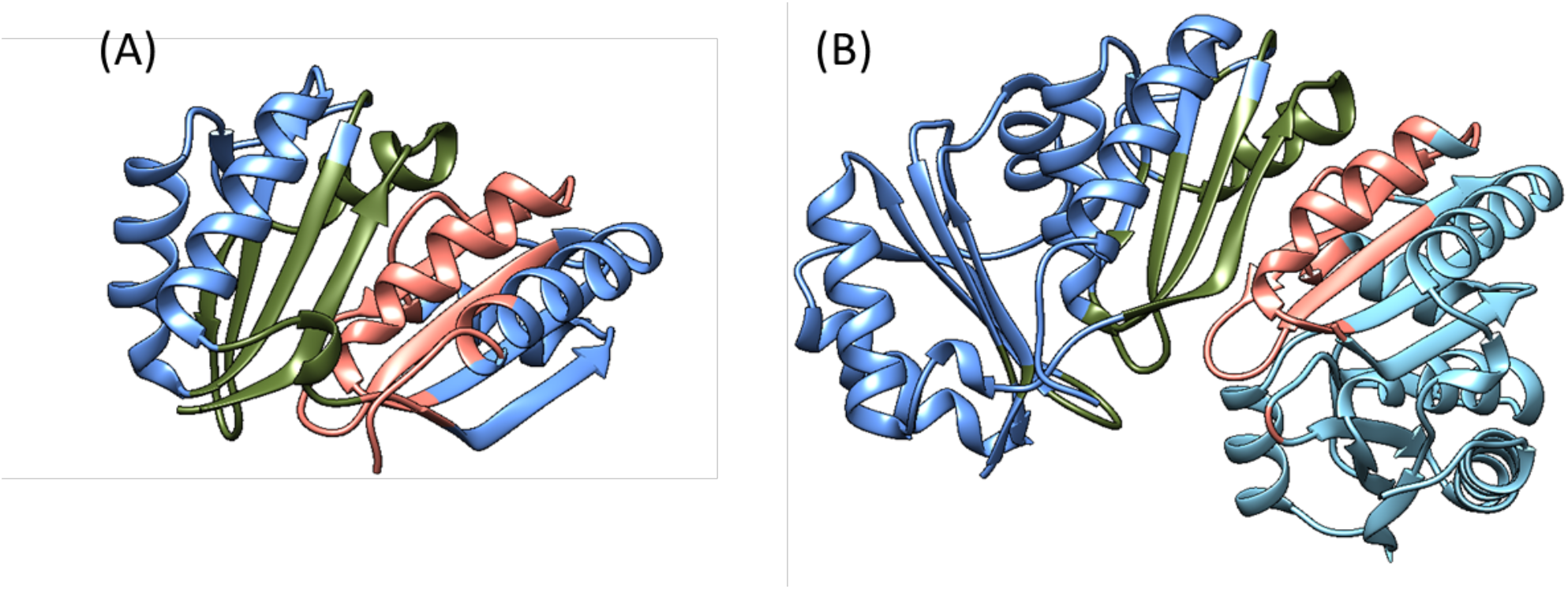
Similarity of chain-chain and domain-domain interfaces. (A) Domain-domain interface between the two domains of carboxysome shell protein CcmP shown in olive green and salmon (B) Chain-chain interface between two chains of carboxysome shell protein CcmP. Each chain is shown in different shades of blue and the interface in olive green and salmon.

#### 2.3.3. Sequence conservation at the interface compared to that of the whole protein

In this section we are comparing the conservation of residues on the interface to the conservation of residues throughout the protein. The sequence identity of the interface was computed as the number of identical residues between the structurally aligned positions of the cluster representative and the members of the cluster. The structural comparison was done using the structural alignment tool CLICK [56]. The sequence identity of the full protein was computed as the number of identical residues computed from a BLAST2seq [57] alignment of the cluster representative with the members of the cluster. Structural alignment was not used while computing the identity of the full protein as the proteins might be structurally dissimilar even though they have structurally similar interfaces. Around 44% of the cases had the interface more conserved, whereas around 37% of the cases had the protein with higher conservation. In 19% of the cases, the interface and the rest of the protein were conserved to the same extent (Figure S8). We investigated the extreme cases where the difference in the identities at the interface and the full protein sequence was > 30%. Most of these were a consequence of non-significant short sequence alignments, non-rigidity of the interface, structurally similar chain-chain and domain-domain interfaces in the same protein and structurally similar chains in heterooligomer (Supplementary Text 2).

##### Clustering of sequentially unrelated interfaces

The interface library shows how some sequentially unrelated/distantly related proteins could have similar interface structures. This library, can in turn, be useful to identify templates (based on structural similarity) to model protein-protein complexes or build structures of multidomain proteins. Around 2% of the interfaces clustered together have an identity of <30% while 10% of them have an identity of <40%. L2-Haloacid dehalogenase from X. autotrophicus (Figure S9A) (PDB ID – 1QQ5) and the hypothetical 2 haloalonoic acid dehalogenase S. tokodaii (Figure S9A) (PDB ID – 2w43) contains the 1.10.150-3.40.50 fold combination. The two proteins are 31% identical to each other. However, the two interfaces superimpose on each other with a structure overlap of 89% and RMSD of 1.47 Å. The Human Psoriasin (Figure S9B) and the Bovine protein SC0067 contain the 1.10.238 fold but are only 27% identical to each other. They have a similar structure and interact using the same geometry with the interface structure overlap of 94.6% and RMSD of 1.0 Å.

### 2.4. Structurally similar protein-protein interfaces from different folds

#### A fold interacting with different folds using the same geometry

The clustering of the interface library was limited to proteins belonging to the same fold combinations, as an all against all comparison of all the interfaces irrespective of their folds is computationally expensive and out of the scope of our computational resources. However, we compared a few interfaces across different folds to check if there exists structural similarity of the interface irrespective of the fold the protein chains/domain belong to.

The NusG (Transcription antitermination protein) and the Transcription elongation factor SPT5 belong to the same fold of alpha-beta plaits and are 34% sequentially identical to each other. The NusG protein interacts with DNA dependent RNA polymerase E, which belongs to Ruberythrin Domain 2-fold whereas the SPT5 interacts with the Transcription elongation factor SPT4 belongs to Herpes Virus 1 fold. Even though, the interacting proteins (DNA dependent RNA polymerase and SPT4) to the two proteins (NusG and SPT5 respectively) belong to different folds and are only 29% identical sequentially the interacting interface is similar with a structure overlap of 93% and RMSD of 1.72 Å (Figure 2). We also found an example of interfaces belonging to different fold that share the same geometry which has been shown in Supplementary Text 3.

**Figure 2.**
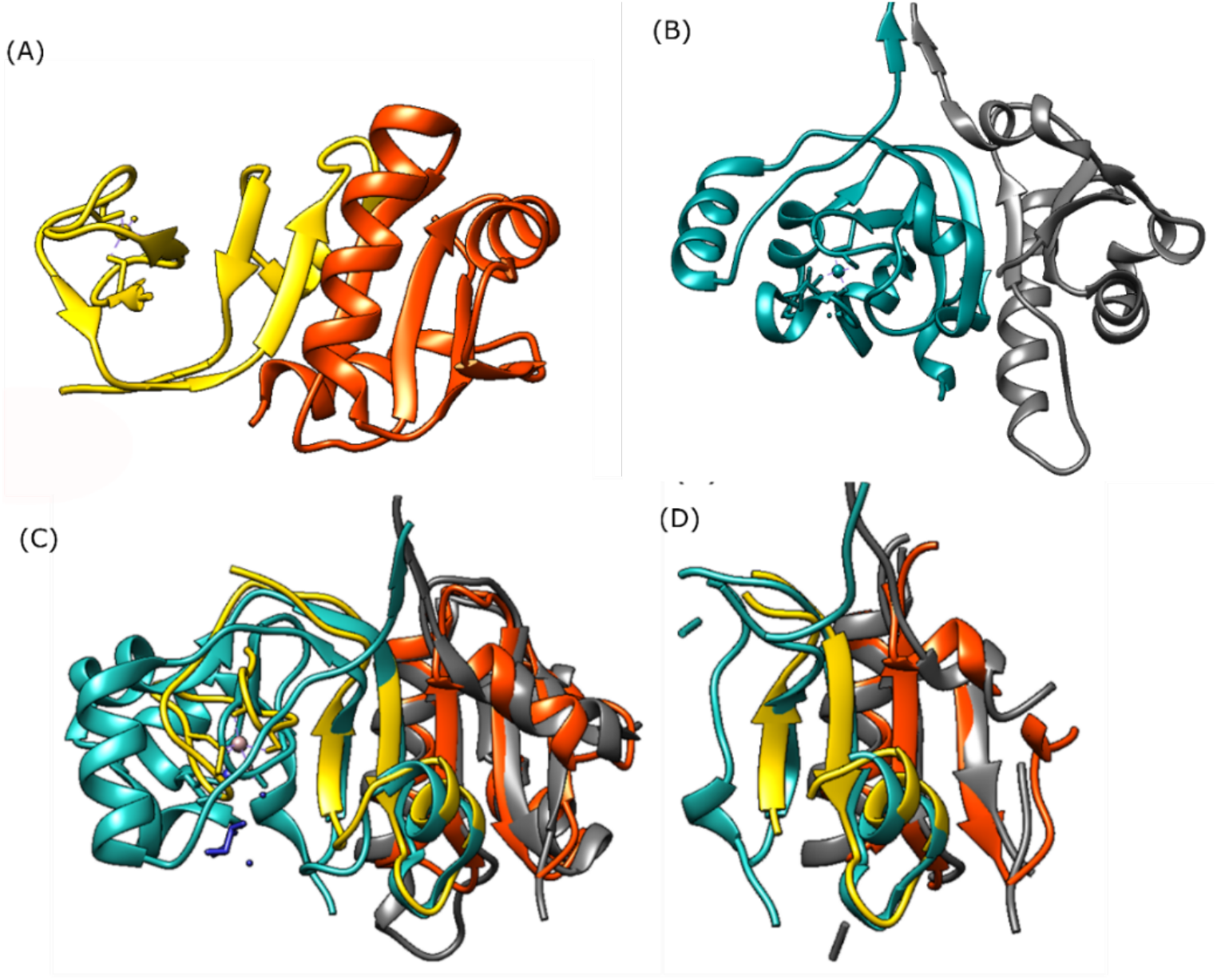
(A) Complex of Nus G protein (PDB – 3LPE_G) (in orange ribbons) and DNA dependent RNA polymerase E (PDB – 3LPE_H) (in yellow ribbons) (B) Complex of SPT5 (PDB – 3H7H_B) (in grey ribbons) and SPT5 and SPT4 (PDB – 3H7H_A) (in cyan ribbons) (C) Superimposition of the complex of Nus G protein and DNA dependent RNA polymerase E onto the complex of (D) The interface residues from the same complexes following the same color scheme shows the structural similarity of the interface residues.

We believe that while we have not systematically categorized such similarities across fold combinations, these data could easily be mined from our database for individual fold combinations.

### 2.5. Pair preference of the amino acid residues at chain-chain vs domain-domain interfaces

The overall trends for the amino acid pair preferences at a chain-chain interface and domain-domain interface look similar (Figure S17). Here, favorable scores are negative numbers, and unfavorable scores are positive. The side chain interactions between Ala with all other amino acids are unfavorable (except Trp, Tyr, Phe, which though favorable, have low scores). Another small amino acid, Cys, also has few favorable interactions. Except for the favorable Cys-Cys interaction (mostly disulphide bridges), its only other favorable interactions (for Met, Asn, Tyr, Trp and Phe) all have low scores. The interactions between the aromatic Tyr, Trp and Phe with all amino acids are favorable at both domain-domain and chain-chain interfaces. However, the interactions of other hydrophobic amino acids - Val, Ile and Leu are usually unfavorable (except low favorable scores with Tyr, Trp and Phe). The interactions between negatively charged amino acids Asp and Glu and positively charged amino acids such as His, Lys, Arg, Gln are favorable, as should be expected. Pi-pi interactions (sp^2^ containing side chains) are found among Tyr, Trp, Phe, His, Arg, Asp, Glu, Asn and Gln [58]. Most of these pairs show high favorability of interaction (potential of >1) except Phe-Asp, Phe-Glu, Asp-Asp, Asp-Glu, Glu-Glu pairs indicating the probable preference for pi-pi interactions at the interface. Self-pairs are preferred in chain-chain interfaces as compared to domain-domain interfaces. This can be because ∼62% of the chain-chain interfaces are homo-oligomers as compared to ∼28% of the domain-domain interfaces being homo-oligomeric (Results Section 2.2).

### 2.6. Small molecule binding site at protein-protein interfaces

Small molecule ligands are often sought to prevent protein-protein interactions [59,60] and have some significance as a therapeutic discovery strategy. Hence, the prediction of small molecule binding sites at the protein-protein interfaces can be the first step toward inhibiting protein complex formation.

The overlap of the residues constituting the binding site and the interface was calculated (Figure S10). Out of the 112,043 chain-chain interfaces, 74,849 interfaces had at least one chain with a minimum of 30% overlap between the predicted binding site and the interface residue. The predicted binding site (using DEPTH) could be used to dock/predict small molecules that could bind at the interface, hence disrupting the formation of the complex [61–64].

One example is that of the inhibition by a small molecule of the interaction between XIAP protein and caspase-9, which is a caspase involved in mitochondrial cell death [65] (Figure 3) [66]. The binding site on the Nipah virus glycoprotein had a 30% overlap with that of the interface residues with the ephrin B2 receptor of humans (PDB – 2VSM). Autodock [67] and DOCK [68] were used to predict the small drug like molecule (ZINC63411510) that would go and bind the predicted binding site of the glycoprotein, hence preventing its interactions with the ephrin-B2 receptor [69].

**Figure 3.**
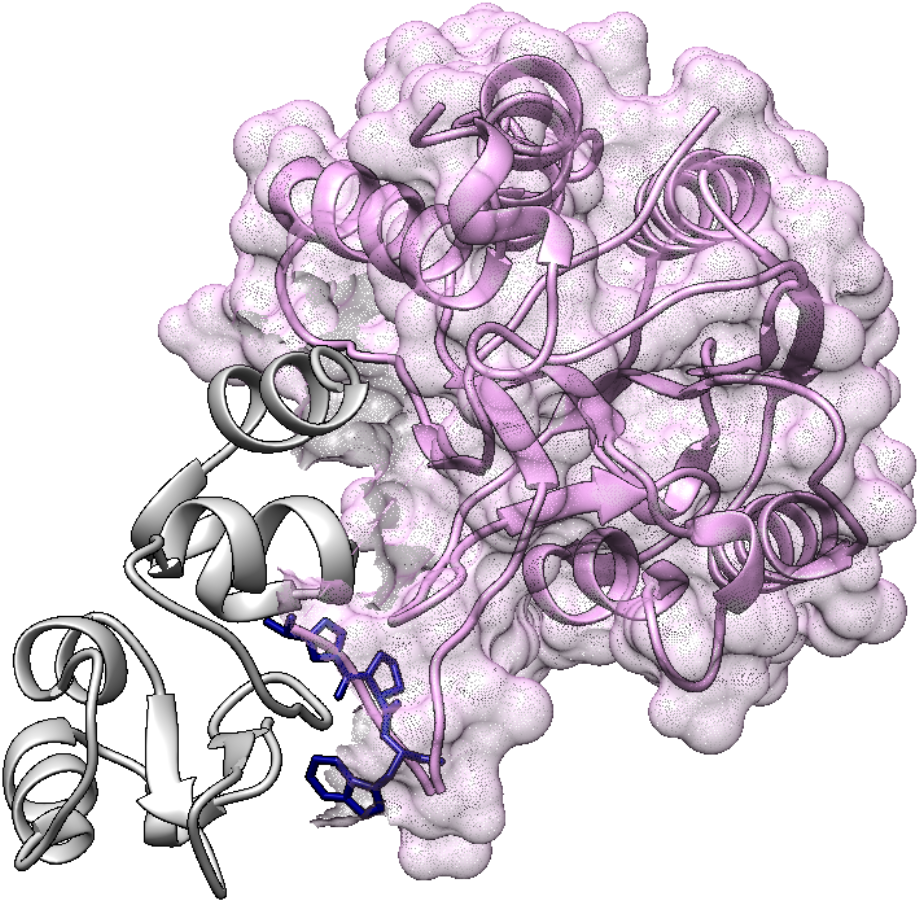
Superimposition of the complex of BIR3 domain of XIAP protein with Caspase 9 (PDB ID – 1NW9) onto the complex of BIR3 domain of XIAP protein with small molecule ligand CO9. CO9 (shown in blue sticks) binds to the region of interaction between BIR3-XIAP (shown in grey ribbons) and Caspase-9 (shown in plum colored ribbon and surface).

## 3. Discussions

Typically, interface libraries contain known associations between fold types. We believe that these associations should be viewed at the level of domains, which could be thought of as a unit of evolutionary conservation. Ideally, we want to annotate interaction patches on domains, but we found that in some complexes where 2 chains interact with one another multiple domains (more than 2) are involved. In such cases, we are unable to determine if the constituent binary domain associations could interact in isolation. Hence, we have organized the data into chain-chain and domain-domain interfaces. Our interface library consists of 112,043 pairs of interacting protein chains and 66,442 pairs of interacting domains taken from crystal structures deposited in the PDB.

Binary chain-chain interactions are typically dominated by homomeric interactions, where the interacting partners have the same CATH id. In our library, 62% of all chain-chain pairs and 28% of domain-domain interactions are homomeric. We conjecture that the other 72% of domain-domain interactions provide us with plausible templates for different types of interactions, many of which have not yet been deposited in the PDB.

In addition to its utility as a plausible template library for modeling protein-protein interactions, our library provides us with several pieces of useful data. The PDB contains 1,391 different CATH folds. Of these, 1109 folds have interacting partners either with itself or another fold. 15 of these folds interact with over 30 other folds. The Rossmann fold, for instance, interacts with 238 other folds. In general, there is a strong correlation (correlation coefficient > 0.8) between the number of EC terms associated with a fold and the number of folds it interacts with. A mismatch between the number of EC terms related to a fold and its number of known interacting partners gives one the basis to search for plausible new interacting partners or EC terms.

One reason for a high number of interfaces for certain folds (folds such as Rossmann fold, Immunoglobulin fold, α helices fold has >1000 interfaces) could be because of the large number of non-homologous proteins that populate these folds (maybe because of convergent evolution). It could also be because of the homo-oligomeric nature of certain folds. Of the 282 folds for which we have no evidence (yet) of interaction with other folds. ∼67% of them belong to the orthogonal bundle (CATH ID – 1.10), irregular architecture (CATH ID – 4.10), 2-layer sandwich (CATH ID – 3.10), alpha-beta complex (CATH ID – 3.90) and up-down bundle (CATH ID -1.20) architecture. Though we cannot conclude this based on the data, we speculate that proteins belonging to these superfamilies do not interact with proteins of other folds.

This library now gives us a platform for examining the nature of interactions between one-fold and its multiple partners. Do date and party hubs use different types of interfaces [70], how could we categorize these, etc. We also examined the extent to which domains and chains use the same interfaces.

Our library has catalogued 3065 fold combinations involving 1109 folds of a possible 1391 folds. Speculatively, even if we assumed that only these 1109 folds were capable of interactions, we could have as many as 614,386 (966,745 fold combinations involving 1391 folds) fold combinations. Clearly, there is a significant mismatch between what is possible and what has been observed, PDB sampling bias and under-representation of complex structures notwithstanding. The extent of the mismatch implies that not all fold combinations are observed in nature. A close examination and analysis of the domain-domain associations in our library may be useful in guiding the construction of interactomes/networks.

The 155,375 interfaces (88,933 chain-chain interfaces+66,442 domain-domain interfaces) with an assigned domain definition clustered into 27,885 clusters based on the structural similarity of the interface. 99.5% of these clusters contained interfaces belonging to the same superfamily within a fold. If the clustering conditions were to be relaxed, more of the interface clusters would contain members from different superfamilies. This indicates that structurally similar interfaces will mostly be found within a superfamily. In turn, this could assist in identifying interfaces on proteins within a superfamily. These interfaces could be homo- or heteromeric. Some of the often-recurring folds such as the Rossmann fold, TIM barrel and Immunoglobulin folds have >900 clusters of homomeric and heteromeric interfaces. This indicates the richness (diversity of sequences taking on the fold) of the fold in the way it explores interaction diversity, which in turn explains the high correlation with EC terms.

100 clusters had both chain-chain and domain-domain interfaces together, irrespective of the sequence similarity. This can indicate gene fusion leading to the formation of a domain-domain interface from a chain-chain interface or gene splitting leading to the formation of a chain-chain interface from the domain-domain interface. An interesting example is that of the CcmP protein, which has structurally similar chain-chain and domain-domain interfaces. Because domain-domain interfaces and chain-chain interfaces are sometimes structurally similar, our library can provide an increased number of templates to model multi-domain proteins whose individual domains may have been crystallized separately.

We observed that the interfaces have higher sequence similarity as compared to that of the whole protein. The sequence similarity and structure similarity go hand and hand however exceptions are noted. We noticed that ∼2% of the interfaces which were clustered together (same fold combination) were structurally similar (structure overlap>80% and RMSD<1.5) but were not sequentially similar (<30% identity). These show that irrespective of the sequence identity this library can be used to search templates for protein complex modeling.

As stated earlier, our primary intent in creating the library was to accumulate a large number of interface templates to model protein-protein interactions. Though we have not explicitly done so in this study, our database when combined with a structure comparison tool, such as CLICK [56], could help us see similarities between interfaces from unrelated folds. Previous studies have also reported how dissimilar folds can use the same geometry at the interface to interact with a certain protein fold. Our database also contains many instances of homologous proteins interacting with each other using structurally different interfaces, such as in lectins, bacterial chemotaxis proteins, ASPP proteins etc. Previous studies have also pointed towards proteins utilizing similar geometry at the interfaces conjecturing that the structural repertoire of interfaces is close to complete [49]. Hence, the interface library presented in this study can serve as a useful resource to model protein complexes using a topology independent structural match to identify templates interfaces for the same.

We also computed the amino acid pair preference at chain-chain and domain-domain interfaces to calculate amino acid substitution scores. Overall, the trends of what amino acid pairs are favored/unfavored are similar in both the chain-chain/domain-domain interfaces. However self-amino acid pairs were generally favored at the chain-chain interface to a greater degree when compared to domain-domain interfaces. This could be because ∼62% of chain-chain interfaces are homo-oligomers whereas only ∼28% of the domain-domain interfaces are homo-oligomers. The similarity between the two pair preferences shows that domain-domain interfaces could supplement chain-chain interfaces and aid in the study of protein-protein interactions.

A significant predictive aspect of our library is the detection of plausible small molecule binding sites on interfaces. About two thirds (67%) of protein-protein interfaces had at least 30% overlap between the interface residues and predicted small molecule binding site residues. This could serve as a useful resource in studying/inhibiting interactions with possible therapeutic applications. With these data, we could possibly analyze the interface structures to determine the most appropriate small molecule that could affect a known interaction.

In this study, we have laid the foundation for future protein-protein and domain-domain interactions studies/predictions/design. The interfaces here could prove useful in constructing the structures of protein complexes or even building a whole protein structure from individually solved domains. Our library could also be used in conjugation with fragment based interface design algorithms such as nanohedra [71]. With the rapid growth of structures resolved using cryo-EM, a library such as ours could also prove useful in refining such structures.

## 4. Methods

### 4.1. Database of interfaces

All multi-chain and multi-domain (based on CATHv4.2 domain definition) complexes were extracted from the PDB. The accessible surface area for the individual protein chains/domains and all possible binary protein chain/domain complexes were calculated using MODELLER [72]. Only binary complexes with greater than 400 Å^2^ change in solvent accessible surface area after interface formation were retained. This cut off was used to filter crystallographic artefacts from biologically relevant interfaces in line with the PQS server [73–76]. Despite this cut-off, some of the interfaces could be artefacts of crystallization. However, we believe that these could still serve as viable templates and the scoring scheme employed for modeling could discern between actual interactions and artefacts [77].

Our library contains a list of interacting amino acid residues. Interacting residues are those that have at least one atom within 8 Å of another atom from a different chain/domain. These interacting residues constitute the interacting interface. The interface library contains dimeric chain-chain or domain-domain interfaces (domain-domain interfaces were between residues of the same chain). Oligomeric interfaces, involving more than 2 chains, are represented in the database by the constituent dimeric interfaces (subject to the same selection described above). All interfaces are also labelled by the CATH folds of their constituent chains/domains and are referred to as a ‘fold combination’ in this study.

### 4.2. Clustering of interfaces

All interfaces with the same fold combination were hierarchically clustered together such that the representative interface is the one with the highest resolution. All interfaces within a cluster were compared to the representative interface using CLICK [78] (a topology independent structural superimposition tool) with C^α^ and C^β^ as representatives atoms for superimposition. An interface was clustered with the representative interface if the structure overlap was >80% and RMSD was <1.5 Å (these values were empirically chosen). A new representative interface was chosen from the remaining interfaces and the same procedure was repeated to find plausible clusters for all interfaces (Figure S11).

21 fold combinations with more than 1000 instances in PDBs were broken into two smaller sets of a maximum of 600 interfaces each (empirically chosen to ensure that the smaller subset had at least 400 members). This reduced the number of structural comparisons for clustering.

### 4.3. Pair preference of amino acids at domain-domain and chain-chain interfaces

A residue-residue interaction profile was calculated for all side chain-side chain interactions using the same statistical potential as described in the method PIZSA [79,80]. The scoring scheme is the ratio of the observed probability to the expected probability of an interface residue pair. To prevent the overrepresentation of certain sequences, the PDB entries were culled using PISCES [81] such that the maximum sequence identity was 40%.

### 4.4 Prediction of small molecule binding sites

The small molecule binding site of the individual chains that form the protein-protein interfaces was predicted with the software DEPTH [82] using default options. DEPTH was earlier compared to other state of art binding site prediction software such as MetaPocket2.0 [83] and Concavity [84] and was shown to be better or at par with these methods [82].

## 5. Data availability

The interface library can be downloaded from <u>here</u>.

## Supplementary Material Description

Interface_library_Supplementary_Information.docx – File containing all the supplementary figures and texts.

## Acknowledgement

MSM would like to acknowledge Wellcome trust-DBT India alliance for senior fellowship. NS would like to acknowledge CSIR-SPMF fellowship. The authors would like to thank COSPI lab members for insightful discussions.

## Notes

https://www.iiserpune.ac.in/~madhusudhan/Interface_library/interface_library.tsv.gz

### Competing Interest Statement

The authors have declared no competing interest.

