## Supplementary File for "A structural database of chain-chain and domain-domain interfaces of proteins"

[*]Corresponding author

#### Supplementary Information

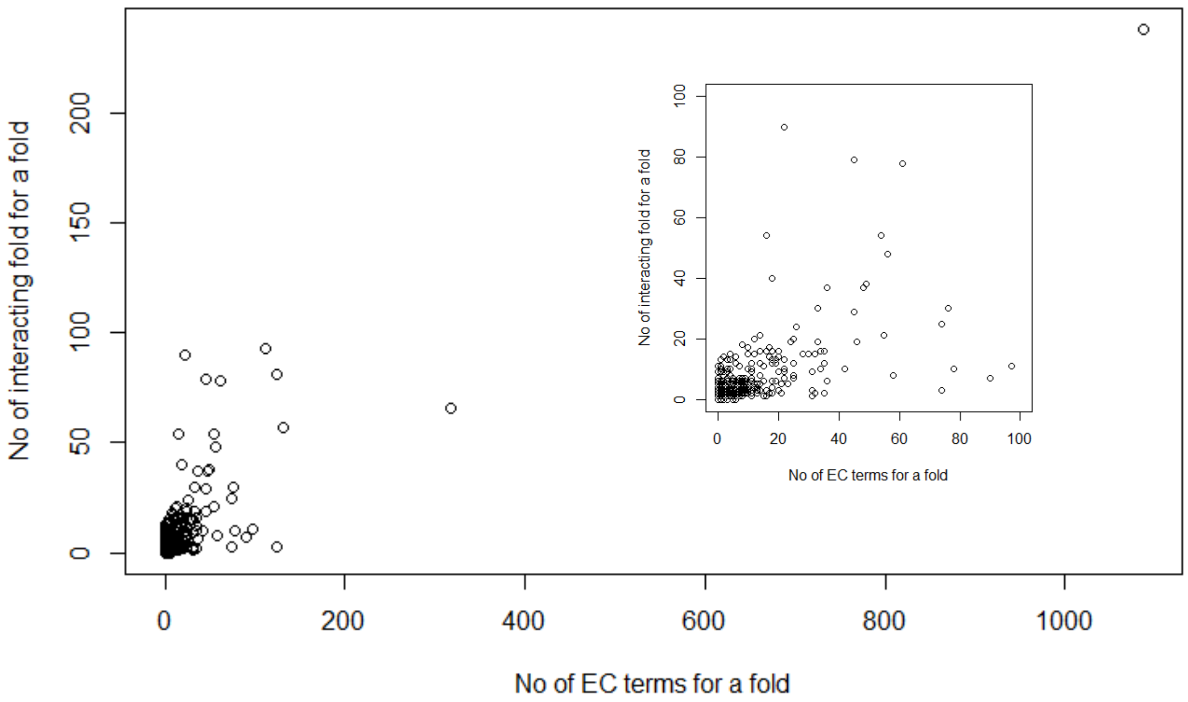


*Figure S1 – The number of EC terms for a fold versus the number of interacting folds for that particular fold. The plot within is the same, except the x and y-axises are truncated at 100.*

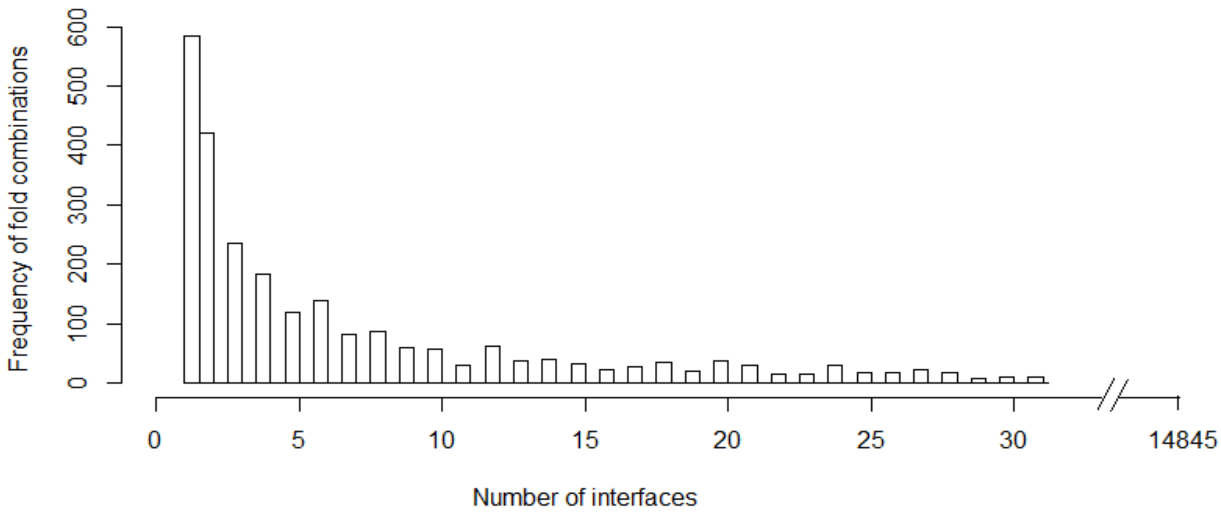


*Figure S2 – Histogram showing the number of fold combinations (y-axis) having a particular number of interfaces (x-axis).*

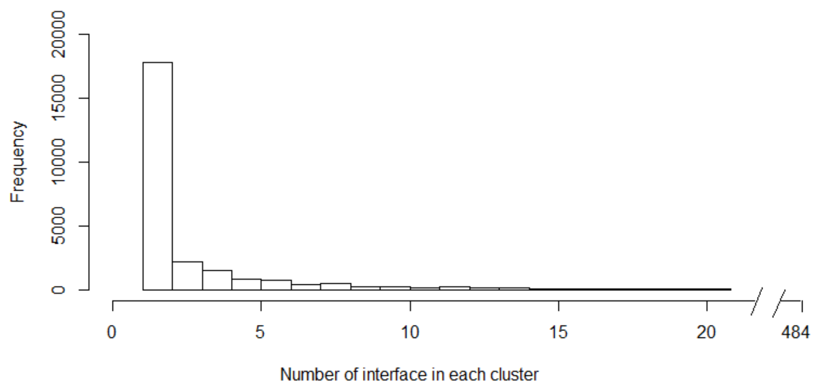


*Figure S3 – Histogram showing the frequency of clusters (y-axis) having a particular number of interfaces in each cluster (x-axis).*

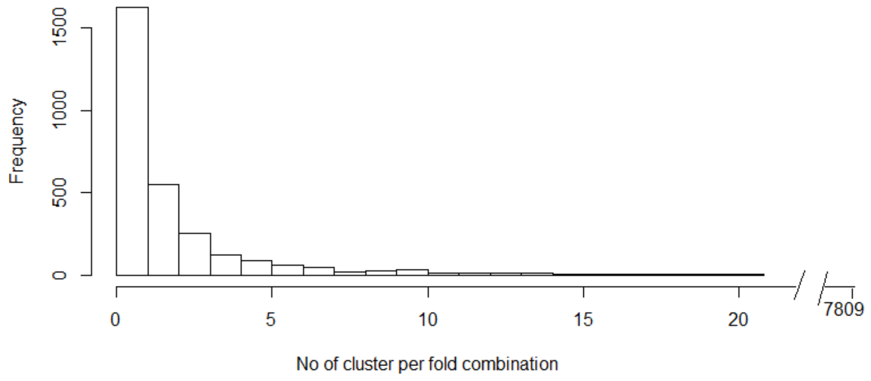


*Figure S4 – Histogram showing the frequency of the number of clusters per fold combination.*

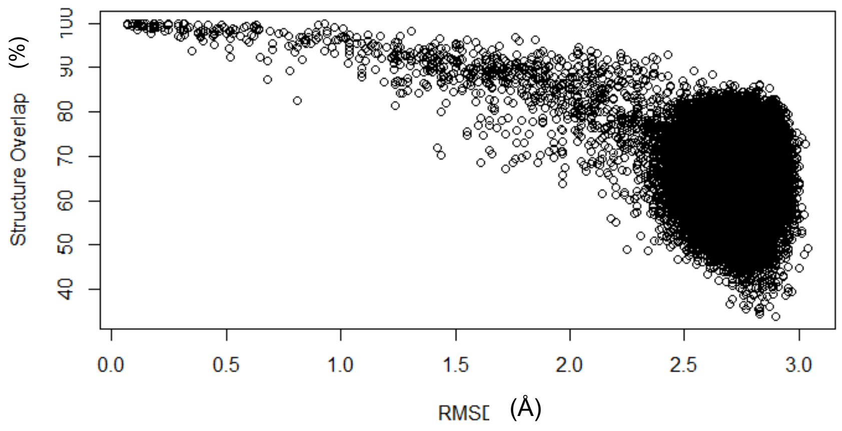


*Figure S5 – Structure overlap (%) vs RMSD (Å) of the structural match between different interfaces belonging to the Rossmann fold.*

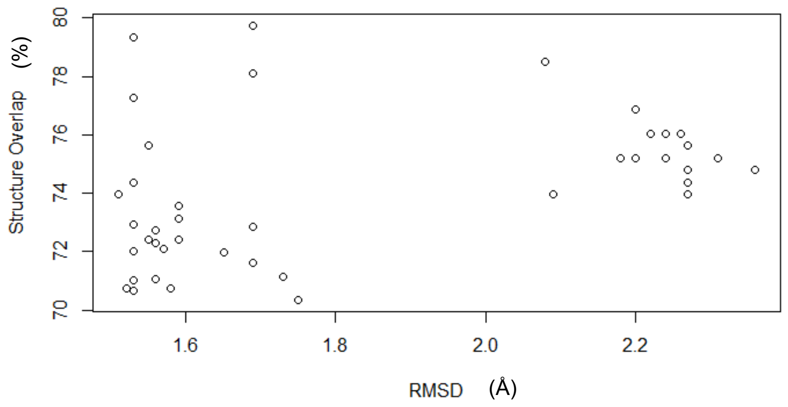


*Figure S6 – RMSD vs Structure Overlap of the interfaces that were not put in the same cluster as the cluster representative 4gxx_BD.*

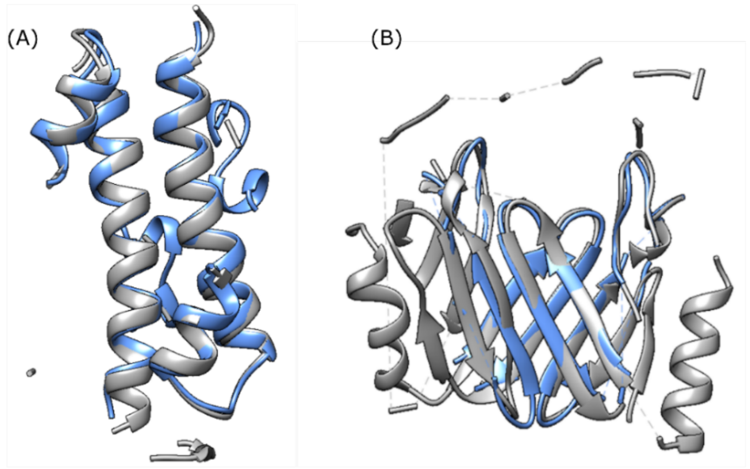


*Figure S7 – (A) Chain-chain interface of Nuclease domain of ribonuclease3 (PDB – 3o2r_CD) in grey ribbons superimposed on the domain-domain interface of Giardia dicer (PDB – 2qvw_B) in blue ribbons (B) Chain-chain interface of AVA_4353 protein (PDB – 3FM2_AB) shown in blue ribbons superimposed on the domain-domain interface of PhuS protein (PDB – 4IMH_B) shown in grey ribbons.*

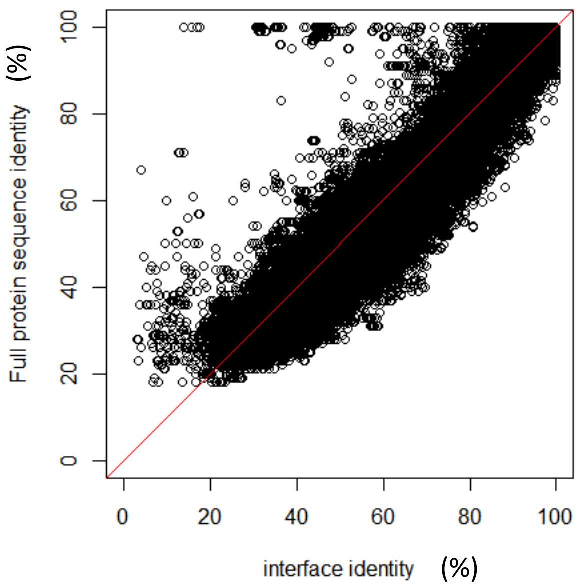


*Figure S8 – Conservation of residues at the interface as calculated from the CLICK structural alignment vs identity of residues when the entire protein is aligned as calculated using BLAST.*

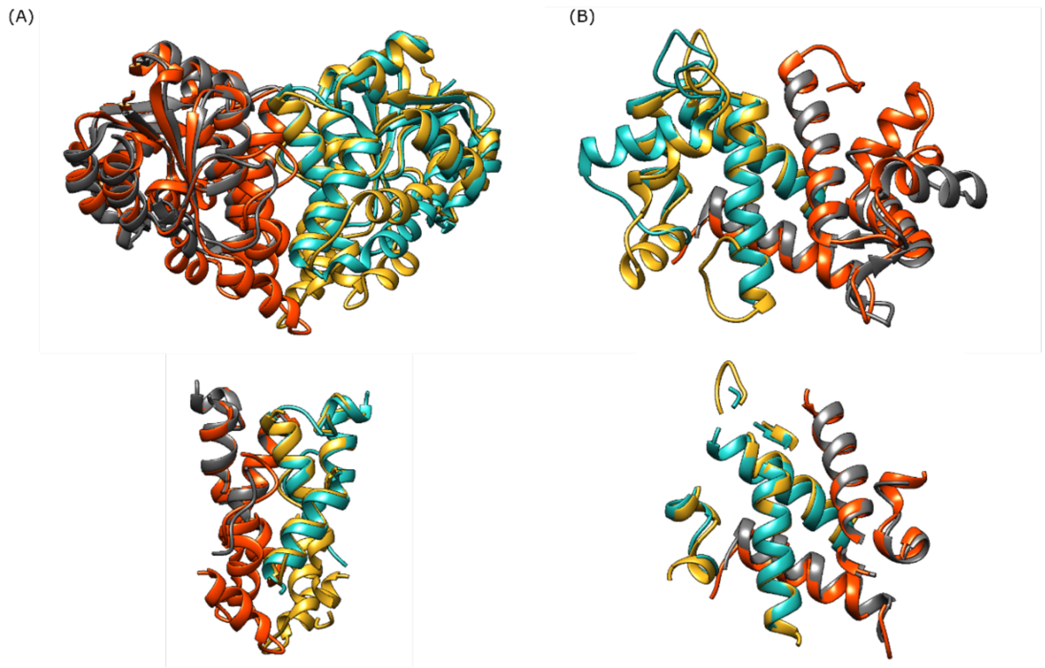


*Figure S9 – (A) The superimposition between PDB 1QQ5_AB (individual chain shown in orange and golden ribbon) and PDB 2w43_AB (individual chain shown in green and grey ribbons). Bottom panel shows the superimposition of the interface residues between PDB*

*1QQ5_AB and 2w43_AB. (B) Top panel shows the superimposition between PDB 1PSR_AB (individual chain shown in orange and golden ribbon) and PDB 3LK0_AB (individual chain shown in green and grey ribbons). Bottom panel shows the superimposition of the interface residues between PDB 1PSQ_AB and 3LK0_AB*

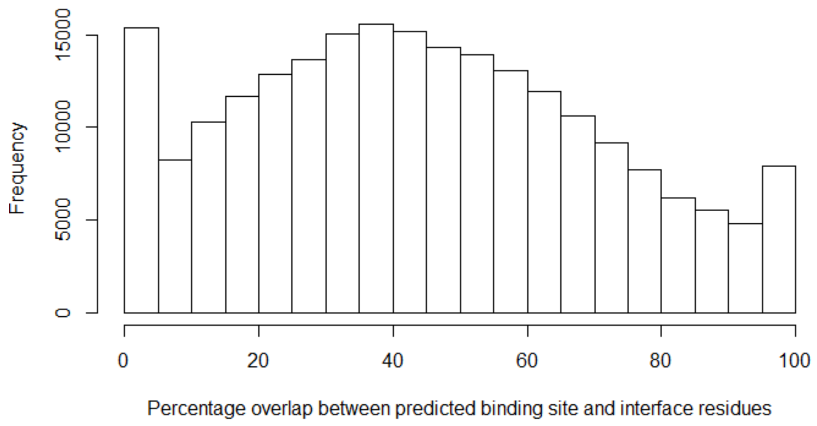


*Figure S10 – Histogram showing the percentage overlap between the predicted small molecule binding sites with that of the interface residues.*

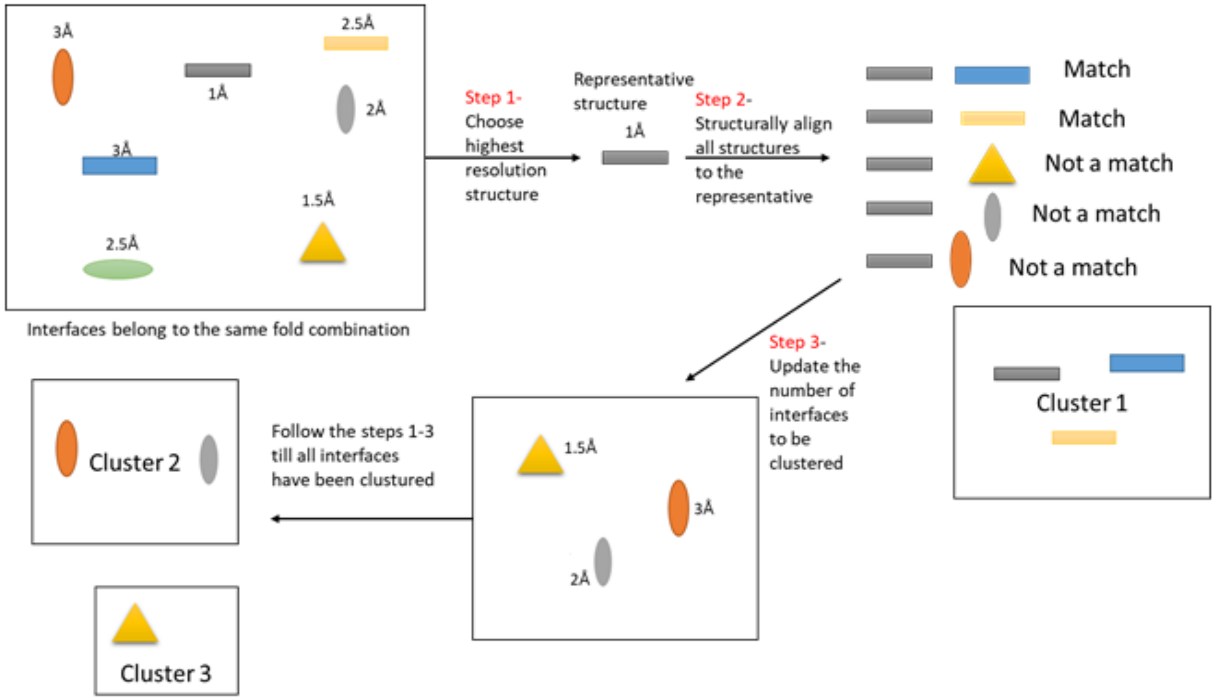


*Figure S11 – Flowchart explaining the clustering of the interface library. The different interfaces belonging to the same fold combination are shown in oval, triangle and rectangle in different colors. The resolution of the structure has been mentioned alongside. The highest resolution structure i.e. grey rectangle is the first representative*

*structure, all other interfaces are structurally aligned to it and considered a match based on a specific criterion. The interfaces, which did not structurally align forms the new set, from which the representative structure is selected, and the steps repeated till all interfaces have been clustered.*

### Supplementary Text

#### Text 1 – Clusters containing both chain-chain and domain-domain interfaces

The clusters containing both chain-chain and domain-domain interfaces varied largely with the number of interfaces varying between 2 to 394 (Figure S12) with a median of 14 interfaces per cluster. For a cluster, the percentage of domain-domain interfaces varied between 0.5%-98.6% with a median of 57% (Figure S13). The percentage of chain-chain interfaces varied between 1.4%-99.5% with a median of 43% (Figure S13).

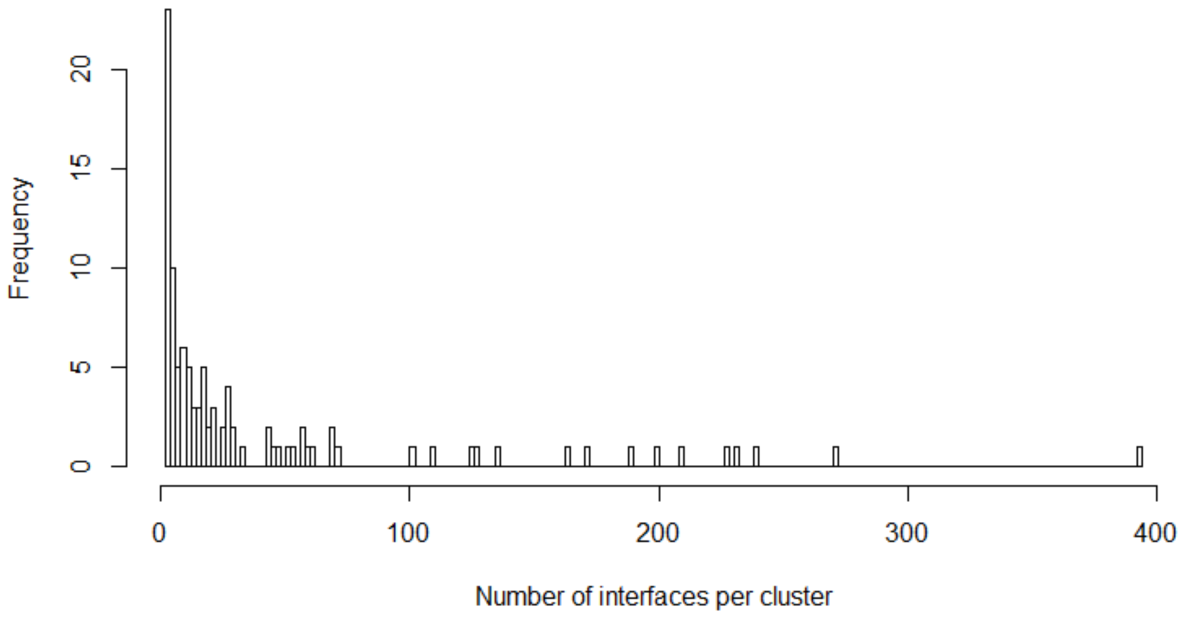


Figure S12 – Frequency of the number of interfaces per cluster where domain-domain and chain-chain interfaces were clustered together.

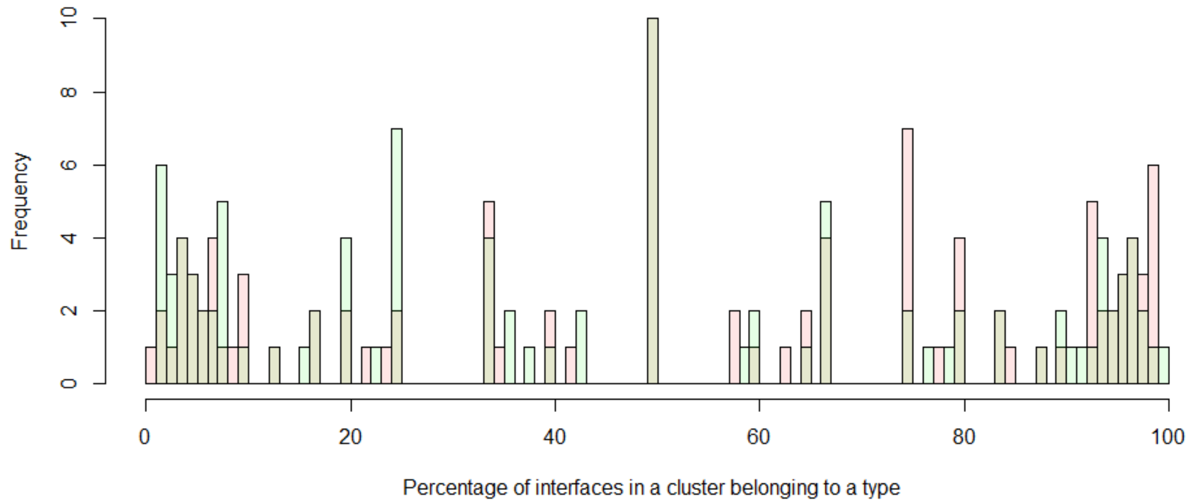


Figure S13 – Frequency of the percentage of interfaces belonging to a particular type (chain-chain/domain-domain). The domain-domain interface frequency has been shown in red, while the chain-chain interfaces has been shown in green.

#### Text 2 – Comparison of the identity of the interface vs that of the full protein

We looked at certain outliers where the difference between the identity of the interface and the full protein was >30%. The cases where the interface identity<20% while the full protein identity is >20% and <80% are because of short local alignments of stretches of ~10 or less residues between the sequences with e-value>1. Hence those alignments were not from significant hits between the protein sequences. The cases where the full protein has ~100% identity while the interface had <40% identity was because of cases where the chain-chain and domain-domain interfaces belonging to the same protein were structurally similar. In these cases, the two domains belonging to the same chain though structurally identical are not sequentially identical. The domain being a part of the chain had 100% identity sequentially, however the identity of the residues at the domain-domain interface and chain-chain interfaces are not same. There are cases where the average sequence identity of the full protein ~100%, however the identity of the interface is between 40-80%. Such cases arise between structurally identical heterooligomer. For example, the Daboitoxin (PDB – 2h4c) is a heterooligomer between Phospholipase A2-III (chain B and F) and Phospholipase A2-II (chain A and E) where the individual monomers have ~60% identity. The two chains are however structurally identical (Figure S14). Hence the structural superimposition of the interface between 2h4c_AB and

2h4c_EF superimposed the A chain of the interface on F chain and B chain on E chain, hence reducing the sequence identity between the interfaces.

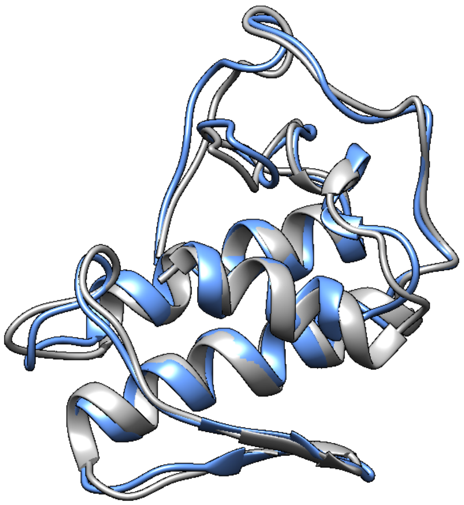

*Figure S14 – Superimposition between the Phosphlipase A2-II (shown in blue ribbons) and Phospholipase A2-III (shown in grey ribbons)*

The cases where the full protein identity is ~100% while the identity of the interface is between 80-90% is because of the flexibility of the whole protein (including the interface) (Figure S15). Also, such cases occur between the alignments of the individual dimeric interfaces of a homo-oligomer (>2 chain). In such case though the individual chains are 100% identical, the residues that form the individual dimeric interfaces vary slightly from one interface to another. Hence the identity of the aligned residues falls below 100%.

In certain cases, such as that of the HTH type transcriptional regulator qacR the two crystal structures (PDB – 3bqz_AB and 1jt6_ED) though sequentially identical, has an RMSD of 0.7 Å (Figure S15). This shows that the whole protein (hence the interface) can be flexible and we can have various forms of the same protein in the PDB. Hence the interface residues and their structural orientation differs between the two PDBs and hence even though the proteins are ~100% identical sequence but has ~90% identity between the aligned interface residues.

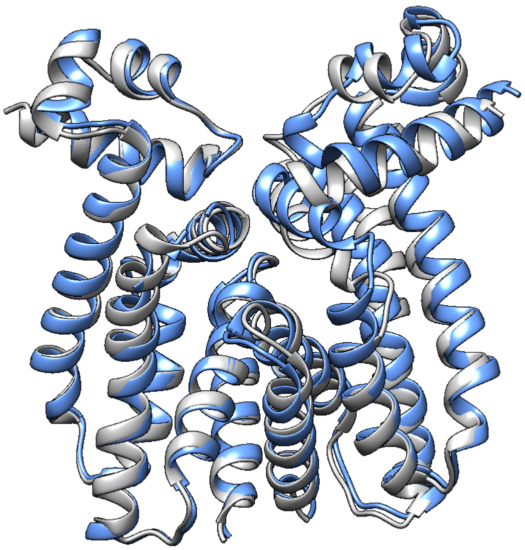

*Figure S15 – Structural superimposition of two different PDBs of HTH type transcriptional regulators qacR (PDB – 3bqz_AB in blue ribbons and PDB – 1jt6_ED in grey ribbons)*

#### Text 3 - Interfaces belonging to different folds utilize the same geometry

The vascular endothelial growth factor A belongs to the cysteine knot cytokine protein whereas the complex between Transcription factor AP1 and c-fos belongs to the single alpha-helices involved in coiled-coil or other helix-helix interface fold. The interacting interface of one involves β-sheets whereas that of others involves α-helices. However, irrespective of the topology of these two unrelated proteins, the two interfaces are structurally similar with a structure overlap of 78% and RMSD of 2.48 Å (Figure S16). We aim to systematically create a database of such cases using our interface library.

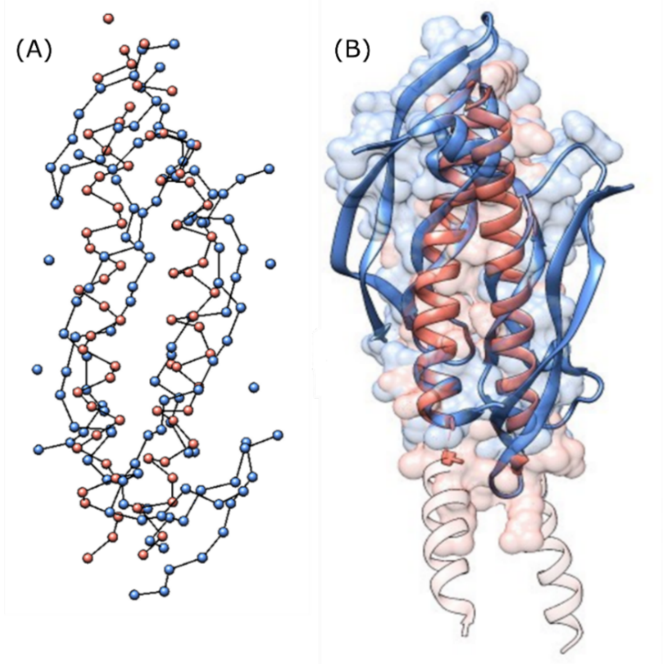


*Figure S16 – (A) Superimposition of the interface of vascular endothelial growth factor A (PDB – 1mkk_AB) (in a blue ball and stick model of the $C^{\alpha}$ atoms) and AP1-c fos (PDB – 1s9k_ED) (in a salmon ball and stick model of the $C^{\alpha}$ atoms) (B) Superimposition of the two protein complexes (shown in ribbons) onto each other following the same color scheme.*

#### Text 4 - Pair preference of the amino acid residues at protein-protein vs domain-domain interfaces

The preference of an amino acid pair to interact with one another in a chain-chain and domain-domain interface was calculated and a statistical potential was computed using the same formulation as PIZSA (Figure S16) at 4 Å distance cut off. The main chain of Gly might form important main chain-main chain interactions or main chain (from Gly)-side chain (from interacting amino acid) interactions. A detailed study of these statistical potentials for chain-chain interfaces at 3 different distance cut-offs of 4 Å, 6 Å and 8 Å for side chain-side chain, side chain-main chain and main chain-main chain interactions can be found at Dhawanjewar *et al.* [80]. However, for this study, we limited ourselves to studying side chain-side chain amino acid pair preferences. Since Gly lacks a side chain, no statistical potential values were computed for pairs containing Gly. Since Gly lacks a side chain, no statistical potential values were computed for pairs containing Gly. For the calculation of the statistical potential for a chain-chain interface a total of 5,571 PDBs (forming 10,836 interfaces) were used, while for the domain-domain interface 2,241 PDBs (forming 2,839 interfaces) were used (sequence being culled at 40% identity).

The pairwise statistical potential for a domain-domain interface ranges between -3.9 to 3.4 whereas that of the chain-chain interface range between -2.1 to 3.8. A negative value indicates that the residue pair is noticed less than expected by random chance (indicating an undesirable interaction) whereas a positive value indicates that the residue pair is noticed greater than by random chance (indicating a favored interaction). The overall trends for the amino acid pair preferences at a chain-chain interface and domain-domain interface look similar (Figure S16), however, a Wilcox paired test show that the two substitution trends are dissimilar with a p-value of 2.2e-16.

(A)

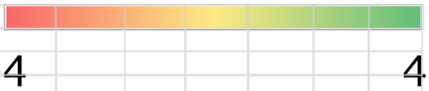


| | ALA | CYS | ASP | GLU | PHE | HIS | ILE | LYS | LEU | MET | ASN | PRO | GLN | ARG | SER | THR | VAL | TRP | TYR |
|---|---|---|---|---|---|---|---|---|---|---|---|---|---|---|---|---|---|---|---|
| ALA | -2.12 | | | | | | | | | | | | | | | | | | |
| CYS | -1 | 3.64 | | | | | | | | | | | | | | | | | |
| ASP | -0.85 | -0.83 | 0.08 | | | | | | | | | | | | | | | | |
| GLU | -1.02 | -0.96 | 0.25 | -0.29 | | | | | | | | | | | | | | | |
| PHE | 0.03 | 1.33 | 0.56 | 0.89 | 2 | | | | | | | | | | | | | | |
| HIS | -1.16 | -0.32 | 2.73 | 2.44 | 1.53 | 2.87 | | | | | | | | | | | | | |
| ILE | -0.52 | 0.07 | -0.54 | -0.35 | 1.36 | 0.52 | 0.52 | | | | | | | | | | | | |
| LYS | -1.94 | -0.44 | 2.67 | 2.42 | 0.43 | 0.62 | -0.06 | -1.11 | | | | | | | | | | | |
| LEU | -1.43 | -0.75 | -1 | -0.75 | 0.76 | -0.04 | 0.44 | -1.1 | -0.77 | | | | | | | | | | |
| MET | -0.29 | 1.16 | -0.11 | 0.43 | 1.58 | 1.09 | 0.9 | 0.23 | 0.18 | 1.69 | | | | | | | | | |
| ASN | -0.78 | 1.51 | 1.86 | 1.72 | 1.24 | 2.46 | 0.55 | 1.31 | -0.11 | 1 | 2.05 | | | | | | | | |
| PRO | -1.37 | -0.48 | 0.1 | 0.43 | 1.09 | 1.15 | -0.06 | -0.38 | -0.54 | 0.21 | 0.73 | -1 | | | | | | | |
| GLN | -0.17 | -0.06 | 1.71 | 1.15 | 1.41 | 1.67 | 0.85 | 1.4 | 0.02 | 1 | 2.17 | 1.15 | 2.72 | | | | | | |
| ARG | -1.56 | -0.17 | 2.95 | 3.01 | 1.28 | 1.07 | -0.31 | -0.56 | -0.74 | -0.03 | 1.59 | 0.04 | 1.77 | 0.29 | | | | | |
| SER | -1.38 | -0.61 | 1.44 | 1.2 | 0.35 | 0.86 | -0.3 | 0.4 | -0.95 | 0.29 | 0.87 | -0.2 | 0.74 | -0.06 | -0.22 | | | | |
| THR | -0.75 | -0.05 | 0.85 | 0.61 | 0.89 | 1.06 | 0.35 | 0.61 | -0.48 | 0.86 | 0.93 | -0.05 | 1.3 | 0.17 | 0.13 | 0.18 | | | |
| VAL | -0.72 | -0.36 | -0.85 | -0.63 | 0.75 | -0.41 | 0.42 | -0.71 | -0.41 | 0.55 | 0.18 | -0.86 | 0.22 | -0.47 | -0.56 | -0.23 | -0.47 | | |
| TRP | 0.42 | 1.48 | 1.68 | 1.79 | 2.73 | 3.28 | 1.99 | 1.79 | 1.08 | 2.46 | 2.16 | 2.57 | 2.16 | 2.21 | 1.27 | 1.73 | 1.38 | 2.64 | |
| TYR | -0.18 | 0.77 | 2.76 | 2.32 | 3.76 | 2.63 | 1.13 | 1.32 | 0.28 | 1.52 | 2.1 | 1.3 | 1.98 | 1.56 | 0.93 | 0.92 | 0.53 | 2.84 | 1.36 |

(B)

| | ALA | CYS | ASP | GLU | PHE | HIS | ILE | LYS | LEU | MET | ASN | PRO | GLN | ARG | SER | THR | VAL | TRP | TYR |
|---|---|---|---|---|---|---|---|---|---|---|---|---|---|---|---|---|---|---|---|
| ALA | -3.92 | | | | | | | | | | | | | | | | | | |
| CYS | -0.43 | 1.3 | | | | | | | | | | | | | | | | | |
| ASP | -0.9 | 0.34 | -1.12 | | | | | | | | | | | | | | | | |
| GLU | -1.23 | -0.34 | 0.18 | -1.55 | | | | | | | | | | | | | | | |
| PHE | 0.14 | 1.39 | 0.78 | 1.08 | 0.17 | | | | | | | | | | | | | | |
| HIS | -0.54 | 1.32 | 2.88 | 2.95 | 1.75 | -0.52 | | | | | | | | | | | | | |
| ILE | -0.22 | 0.9 | -0.16 | 0.36 | 1.56 | 0.94 | -0.87 | | | | | | | | | | | | |
| LYS | -2.13 | -0.47 | 2.86 | 2.77 | 0.38 | 0.99 | 0.09 | -2.61 | | | | | | | | | | | |
| LEU | -0.7 | 0.38 | -0.59 | -0.6 | 0.91 | 0.51 | 1.05 | -0.86 | -1.28 | | | | | | | | | | |
| MET | -0.35 | 1.36 | 0.34 | 0.49 | 1.93 | 1.28 | 1.67 | -0.21 | 1.09 | -0.65 | | | | | | | | | |
| ASN | -0.42 | 0.55 | 2.33 | 1.86 | 1.96 | 1.81 | 0.46 | 1.34 | 0.07 | 1.32 | 0.48 | | | | | | | | |
| PRO | -1.18 | 0.39 | -0.13 | 0.34 | 0.66 | 0.8 | 0.36 | -0.59 | -0.24 | 1.04 | 0.87 | -2.16 | | | | | | | |
| GLN | -0.23 | 1.7 | 1.44 | 1.65 | 1.99 | 2.4 | 0.84 | 1.91 | 0.49 | 1.95 | 2.78 | 1.17 | -0.02 | | | | | | |
| ARG | -1.36 | 0.15 | 3.2 | 3.4 | 1.24 | 1.94 | 0.26 | -0.44 | -0.3 | 0.37 | 1.82 | 0.05 | 1.72 | -1.5 | | | | | |
| SER | -1.26 | 0.09 | 1.93 | 1.86 | 0.53 | 1.02 | 0.02 | 0.7 | -0.61 | 0.3 | 1.58 | -0.19 | 1.44 | 0.13 | -2.18 | | | | |
| THR | -0.45 | 0.22 | 1.31 | 1.37 | 0.96 | 1.24 | 0.38 | 0.11 | 0.16 | 0.76 | 1.67 | 0.27 | 1.47 | 0.46 | 0.53 | -1.85 | | | |
| VAL | -0.81 | 0.47 | -0.66 | -0.34 | 0.88 | 0.16 | 0.55 | -1.04 | 0.04 | 0.42 | 0.28 | -0.24 | 0.54 | -0.49 | -0.69 | 0.02 | -2.08 | | |
| TRP | 0.53 | 2.57 | 2.22 | 2.14 | 2.98 | 3.1 | 2.16 | 2.83 | 1.69 | 3.25 | 2.23 | 2.46 | 2.72 | 2.85 | 1.96 | 2.51 | 1.5 | 1.02 | |
| TYR | 0.04 | 1.72 | 2.85 | 2.69 | 2.1 | 2.51 | 1.11 | 1.51 | 0.68 | 1.95 | 2.12 | 1.48 | 1.84 | 1.72 | 1.06 | 1.1 | 0.74 | 2.97 | -0.26 |

*Figure S17 – Statistical potential for the pairwise interaction between 2 amino acids from different chains for (A) chain-chain (B) domain-domain interfaces. Each of the pair preference is colored based on their pair preference score.*

The difference between the domain-domain score and the chain-chain scores were computed (Figure S18). Self-pairs are seen to be preferred in chain-chain interfaces as compared to domain-domain interfaces. We notice self-amino acid pairing (other than Cys, Asn, Phe and Trp) are not favored at the domain-domain interface i.e. have negative values. However, most self-amino acid pairing other than Ala, Lys, Leu, Phe and Val (Glu and Ser to some extent) are favored at the chain-chain interface. This can predominantly be because ~62% of the chain-chain interfaces are homo-oligomers as compared to ~28% of the domain-domain interfaces being homo-oligomeric (Results Section 2.2). We notice that Cystine interactions in domain-domain interfaces have lesser scores than that of the chain-chain interfaces, probably because of its lower natural abundance, and fewer number of domain-domain interfaces (2839 domain-domain interfaces compared to 10836 chain-chain interfaces).

-2 3

| | ALA | CYS | ASP | GLU | PHE | HIS | ILE | LYS | LEU | MET | ASN | PRO | GLN | ARG | SER | THR | VAL | TRP | TYR |
|---|---|---|---|---|---|---|---|---|---|---|---|---|---|---|---|---|---|---|---|
| ALA | 1.8 | | | | | | | | | | | | | | | | | | |
| CYS | -0.56 | 2.34 | | | | | | | | | | | | | | | | | |
| ASP | 0.04 | -1.2 | 1.2 | | | | | | | | | | | | | | | | |
| GLU | 0.21 | -0.6 | 0.07 | 1.26 | | | | | | | | | | | | | | | |
| PHE | -0.11 | -0.1 | -0.23 | -0.2 | 1.83 | | | | | | | | | | | | | | |
| HIS | -0.62 | -1.6 | -0.15 | -0.5 | -0.22 | 3.38 | | | | | | | | | | | | | |
| ILE | -0.31 | -0.8 | -0.38 | -0.7 | -0.2 | -0.43 | 1.39 | | | | | | | | | | | | |
| LYS | 0.19 | 0.03 | -0.19 | -0.4 | 0.05 | -0.37 | -0.15 | 1.5 | | | | | | | | | | | |
| LEU | -0.73 | -1.1 | -0.41 | -0.2 | -0.15 | -0.55 | -0.62 | -0.23 | 0.51 | | | | | | | | | | |
| MET | 0.06 | -0.2 | -0.45 | -0.1 | -0.35 | -0.19 | -0.77 | 0.44 | -0.91 | 2.34 | | | | | | | | | |
| ASN | -0.36 | 0.96 | -0.47 | -0.1 | -0.72 | 0.64 | 0.09 | -0.03 | -0.18 | -0.32 | 1.57 | | | | | | | | |
| PRO | -0.18 | -0.9 | 0.23 | 0.1 | 0.42 | 0.36 | -0.42 | 0.21 | -0.31 | -0.83 | -0.14 | 1.16 | | | | | | | |
| GLN | 0.06 | -1.8 | 0.27 | -0.5 | -0.58 | -0.72 | 0.01 | -0.51 | -0.47 | -0.94 | -0.61 | -0.03 | 2.75 | | | | | | |
| ARG | -0.21 | -0.3 | -0.25 | -0.4 | 0.04 | -0.87 | -0.57 | -0.12 | -0.44 | -0.4 | -0.23 | -0.01 | 0.05 | 1.79 | | | | | |
| SER | -0.12 | -0.7 | -0.49 | -0.7 | -0.17 | -0.16 | -0.32 | -0.29 | -0.34 | -0.01 | -0.71 | -0.01 | -0.7 | -0.19 | 1.96 | | | | |
| THR | -0.31 | -0.3 | -0.46 | -0.8 | -0.07 | -0.18 | -0.03 | 0.51 | -0.64 | 0.1 | -0.74 | -0.32 | -0.18 | -0.28 | -0.4 | 2.03 | | | |
| VAL | 0.09 | -0.8 | -0.2 | -0.3 | -0.13 | -0.57 | -0.13 | 0.32 | -0.45 | 0.13 | -0.1 | -0.63 | -0.32 | 0.02 | 0.14 | -0.26 | 1.62 | | |
| TRP | -0.11 | -1.1 | -0.54 | -0.4 | -0.26 | 0.18 | -0.18 | -1.04 | -0.61 | -0.79 | -0.07 | 0.11 | -0.56 | -0.64 | -0.69 | -0.78 | -0.12 | 1.62 | |
| TYR | -0.22 | -1 | -0.09 | -0.4 | 1.67 | 0.12 | 0.02 | -0.19 | -0.4 | -0.42 | -0.02 | -0.18 | 0.14 | -0.16 | -0.12 | -0.18 | -0.2 | -0.13 | 1.61 |

*Figure S18 – Difference in amino acid pair potential (Chain chain interface score – Domain domain interface score) colored based on the difference.*

#### Text 5 - Comparison of SCOPe and CATH domain definition

The number of folds in SCOPe (version – SCOPe-2.07-stable) and CATH (version - b.20180915) are 1,457 and 1,391 respectively. Out of these 377 and 282 folds (for SCOPe and CATH respectively) have no other domain interacting partner in the PDB. Only 31,063 PDBs had domain definitions (other than C terminal tag defined by tag l.1.1) according to SCOPe, whereas 114,839 PDBs had domain definitions according to CATH. The number of assigned multidomain PDBs according to SCOP is 17,303 while that according to CATH is 43,784. SCOPe assigns 57 proteins wherein a single domain is defined such that it spans multiple chains, whereas CATH has no such anomalous cases. Out of 112,043 chain-chain interfaces as observed in the PDB, 49,888 chain-chain interfaces did not have a fold assigned to at least 1 chain according to SCOPe definition, while 22,110 chain-chain interfaces did not have a fold classification according to CATH. Hence, because of the larger coverage of the domain definition in CATH for the PDB as compared to SCOPe, the CATH domain definition has been used for the interface library creation.
